# Bacterial lipoproteins induce distinct IL-8 inductions relying on their lipid moieties in pulmonary epithelial cells

**DOI:** 10.1101/2020.04.03.024471

**Authors:** Sang Su Woo

## Abstract

Pulmonary epithelial cells play a crucial role in host defense against bacterial insult. However, the innate immunity in pulmonary epithelium is relatively less characterized than that of professional immune cells. In the current study, we investigated IL-8 induction by bacterial cell wall components in human lung epithelial A549 cells and primary human bronchial epithelial cells. We found that lipoproteins are the most potent IL-8 inducers among bacterial cell wall components. Using synthetic lipopeptides that mimic bacterial lipoprotein pam2CSK4 and pam3CSK4, we further investigated signal transduction mechanism. Interestingly, IL-8 secretion appeared higher upon diacylated pam2CSK4 treatment than triacylated pam3CSK4. A role of TLR2 in lipopeptide-induced IL-8 secretion was monitored by siRNA transfection and functional blocking antibodies. We found that TLR2 in A549 cell localizes to intracellular compartment. Activation mechanism for the intracellular TLR2 differed between pam2CSK4 and pam3CSK4. Pam2CSK4 utilized a lipid-raft dependent mechanism while pam3CSK4 utilizes clathrin-dependnet endocytosis. Using CHO/CD14/TLR2 cell line, we found the distinct IL-8 induction level initiates at the receptor activation level. Subsequent pharmacological inhibitor studies showed that pam2CSK4 and pam3CSK4 utilizes different MAPK pathways. Downstream transcription factor activation was determined by electro-mobility shift assay (EMSA) and luciferase reporter assays. The results showed distinct pattern of activation for NF-κB, NF-IL6 and AP-1 from human IL-8 promoter. These results suggest that bacterial lipoproteins from cell walls are an important determinant in initiating tissue inflammation in pulmonary epithelium upon bacterial insult.

## 1. Introduction

Bacterial infection in airway epithelium plays a critical role in inducing acute and inflammatory diseases such as pneumonia and chronic obstructive pulmonary disease (COPD) and cystic fibrosis (CF) via pattern-recognition receptor activation (PRR) (Dickson, Martinez, & Huffnagle, 2014; Huffnagle & Dickson, 2015; Moldoveanu et al., 2009). The activation of PRRs in airway epithelium subsequently activate resident alveolar macrophages and recruit second line defense mechanism such as neutrophils (Moldoveanu et al., 2009). Neutrophils are recruited to parenchymal tissues following elevated release of chemokine Interleukin-8, which is typically expressed in antigen presenting cells (Yoshimura et al., 1987). The infiltrated neutrophils elicit tissue inflammation and damages (Riches & Martin, 2018). However, direct regulation of lung epithelial expression and release of IL-8 is relatively poorly studied (Wickremasinghe, Thomas, & Friedland, 1999).

Bacterial lipopolysaccharide (LPS) has been considered a major virulence factor that induces immune response against pulmonary pathogens since it systematically activates dendritic cells and macrophages (Eisenbarth et al., 2002; Rodriguez et al., 2003) However, in lung epithelial cells, the corresponding PRR, toll-like receptor 4 (TLR4) has been found in intracellular compartment (Guillot et al., 2004). Therefore, it remained to be addressed whether bacterial pathogen-associated molecular pattern (PAMP) directly induces IL-8 expression in lung epithelial cells. It was reported that bacterial flagellin induces strong IL-8 secretion in human lung epithelial cells (Im et al., 2009). Among other bacterial cell wall-derived PAMPs, lipoteichoic acids (LTA), peptidoglycans (PGN) and lipoproteins were suggested as virulence factors for TLR2 activation in airway and lung (Bubeck Wardenburg, Williams, & Missiakas, 2006; Cheon et al., 2008; Triantafilou et al., 2004), among which bacterial lipoprotein appeared indispensable for *Staphylococcus aureus* infection (Bubeck Wardenburg et al., 2006).

Different bacteria exhibit distinct pattern of lipoproteins. Bacterial lipoproteins are synthesized through operation of three distinct enzymes, lipoprotein diacylglyceryl transferase (Lgt), prolipoprotein signal peptide (Lsp), and apolipoprotein *N*-acyltransferase (Lnt) (Kovacs-Simon, Titball, & Michell, 2011). Lgt lipidates prolipoproteins containing signal peptides and produces diacylated lipoproteins; Lsp cleaves signal sequence; Lnt catalyzes additional acylation to produce triacylated lipoprotein. Gram-negative bacteria typically require all three enzymes for their survival whereas Lsp is dispensable in Gram-positive bacteria. By consequence, Gram-negative bacteria usually maintain triacylated lipoproteins while Gram-positive bacteria including *Staphylococci* such as *S. aureus* possess diacylated lipoproteins (Bubeck Wardenburg et al., 2006). This suggests that different bacterial species present different arrays of lipoproteins. Besides that diacylated lipoproteins bind to TLR2/TLR6 whereas triacylated lipoproteins bind to TLR2/TLR1 (Nakayama, Kurokawa, & Lee, 2012), information regarding direct immune response from pulmonary epithelial cells against bacterial lipoproteins is sparse (Pabst, Durak, Roos, Luhrmann, & Tschernig, 2009; Reppe et al., 2009; Waters et al., 2005).

In the current study, we examined various bacterial cell wall-derived PAMPs on IL-8 secretion in human lung and airway epithelial cells. Among PAMPs tested, *S. aureus* lipoprotein showed the most potent IL-8 induction. We hypothesized that different lipid moieties might be responsible for different IL-8 induction. Using synthetic lipopeptides with different lipid moieties, (pam2CSK4 for diacylated and pam3CSK4 for triacylated), we confirmed that diacylated lipopeptides possess a stronger immunogenicity to induce IL-8 secretion than triacylated lipopetides in human pulmonary epithelial cells. Receptor activation and downstream signal transduction via MAPKs were surveyed using siRNA transfection and pharmacological inhibitors. Differential activation of transcription factors by diacylated and triacylated lipopeptides was also monitored.

## Results

### *S. aureus* bacterial lipopeptide is a potent IL-8 inducer in human lung epithelial cells

We examined cell wall components from Gram-negative bacteria *E. coli* and Gram-positive bacteria *S. aureus* for IL-8 secretion in human type II alveolar epithelial cell, A549 cells. LPS, LTA, PGN and lipoproteins were treated in varying doses for 24 hr and IL-8 secretion was determined by ELISA (Fig. 1A). *S. aureus* lipoprotein exhibited the most potent IL-8 induction with a ng/ml level of IL-8 in culture supernatant in a dose-dependent manner. *S. aureus* LTA and PGNs also induced IL-8 secretion but no more than 0.2 ng/ml. It was evident that neither LPS nor lipoprotein from *E. coli* significantly elicited IL-8 secretion. We further confirmed this finding using EtOH-killed *S. aureus* deficient of either lipoprotein (Δlgt) or LTA (ΔltaS) in their cell wall (A. R. Kim et al., 2017; J. Kim et al., 2013). Both EtOH-killed parental and ΔltaS strains induced 1 ng/ml level IL-8 secretion, whereas Δlgt strain showed a markedly reduced IL-8 secretion comparable to untreated control (Fig. 1B). These results showed that *S. aureus* lipoprotein is a potent bacterial cell wall-derived PAMP that induces IL-8 secretion in human lung epithelium.

**Fig. 1.**
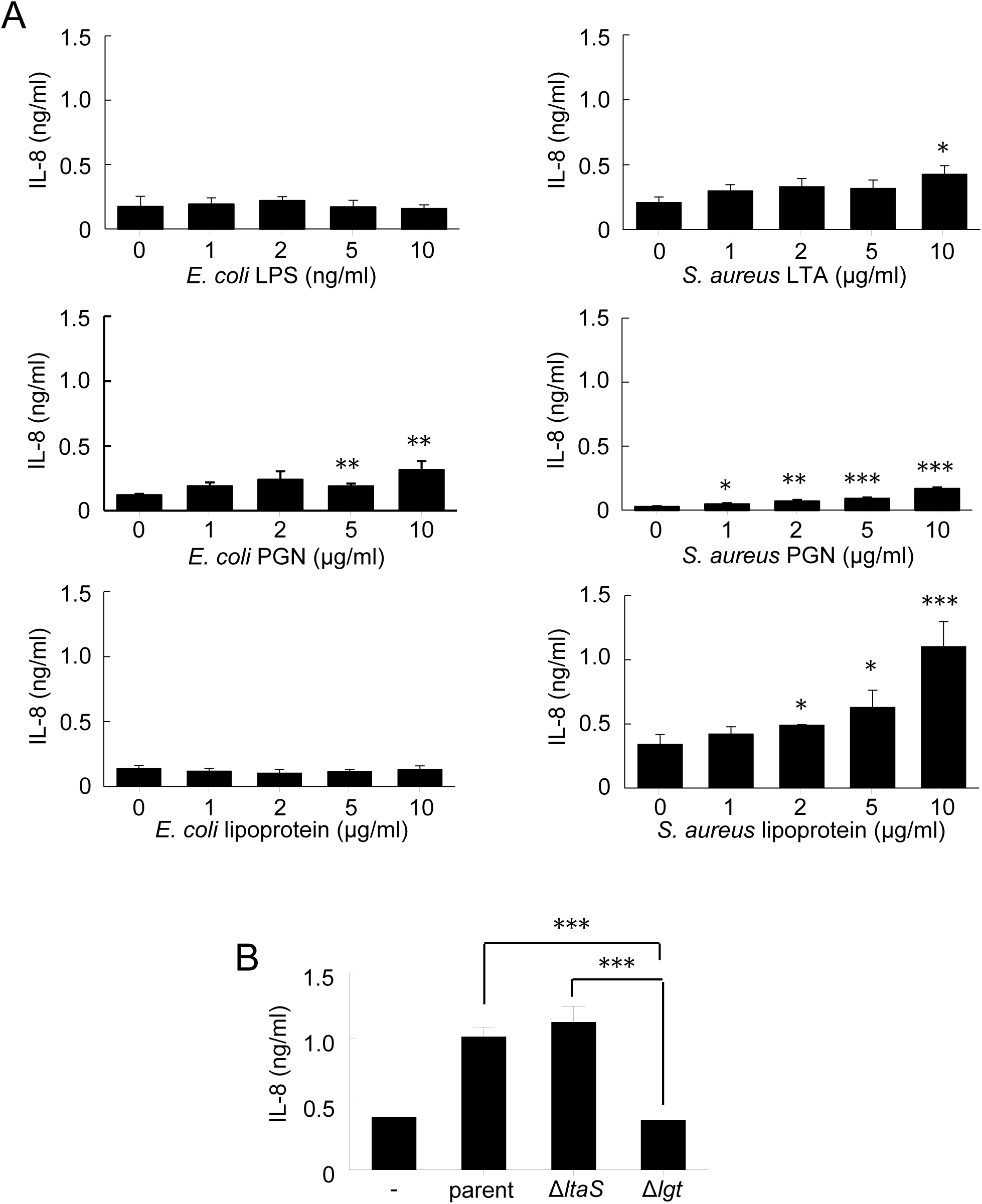
IL-8 induction induced by E. coli and S. aureus cell wall-derived PAMPs. (A) Human lung epithelial A549 cells were treated with indicated amount of bacterial PAMPs for 24 h. IL-8 secretion was determined by ELISA for culture supernatants. Left-side panels show IL-8 secretion upon treatment of *E. coli* cell wall-derived PAMPs. From top to bottom, LPS, PGN or lipoprotein; right-side panels exhibit IL-8 secretion upon *S. aureus* cell wall-derived PAMPs. From top to bottom, LTA, PGN, and lipoprotein. (B) Effect of lipoprotein deficiency in IL-8 production from A549 cells. An ELISA experiment was performed as in A. Cells were treated with 200 μg/ml of ethanol-inactivated *S. aureus* strains: parental, LTA-deficient (ΔltaS) or lipoprotein-deficient (Δlgt) for 24 h. Bars indicate mean ± S.D. (n=3).

### Diacylated lipopeptide is more potent than triacylated lipopeptide for IL-8 induction in pulmonary epithelial cells

In following studies, we used synthetic lipopeptides with different lipid moiety (pam2CSK4, diacylated; pam3CSK4, triacylated). First, we tested whether they induce the release of IL-8 in A549 cells by ELISA (Fig. 2A-B). Although both pam2CSK4 and pam3CSK4 induced ng/ml level of IL-8 secretions, IL-8 secretion in cells treated with pam2CSK4 was 4-fold higher than that of pam3CSK4 at the same lipopeptide concentration. We found this difference interesting and decided to further characterize. The differential activity of different lipopeptides was examined in *ex vivo* condition using primary human bronchial epithelial cells. IL-8 secretion was determined by ELISA for human primary bronchial epithelial cells upon lipopeptide treatment. Indeed, pam2CSK4 was more efficient for IL-8 secretion than pam3CSK4 (Fig. 2B). We also monitored time-dependent IL-8 protein secretion in A549 cells (Fig. 2C). Pam2CSK4-induced IL-8 secretion was higher than pam3CSK4 for 48 hr incubation. This result suggested that pam2CSK4 induces stronger IL-8 mRNA expression than pam3CSK3. Quantitative RT-PCR for IL-8 mRNA validated that pam2CSK4 elicits greater IL-8 mRNA expression than pam3CSK4 (Fig. 2D). Again, this was not due to a specific incubation time point since IL-8 mRNA expression induced by pam2CSK4 was constantly higher than that of pam3CSK3 during 12 hr treatment (Fig. 2E). It was noteworthy that pam2CSK4-induced IL-8 mRNA expression reached its peak at around 6 hr while pam3CSK4-induced IL-8 mRNA express reached its peak at 3 hr. This result suggested that IL-8 induction between pam2CSK4 and pam3CSK4 could result from different signal transduction pathways. Collectively, these data showed that diacylated pam2CSK4 induces higher level of IL-8 secretion than tri-acylated pam3CSK4 in human pulmonary epithelial cells.

**Fig. 2.**
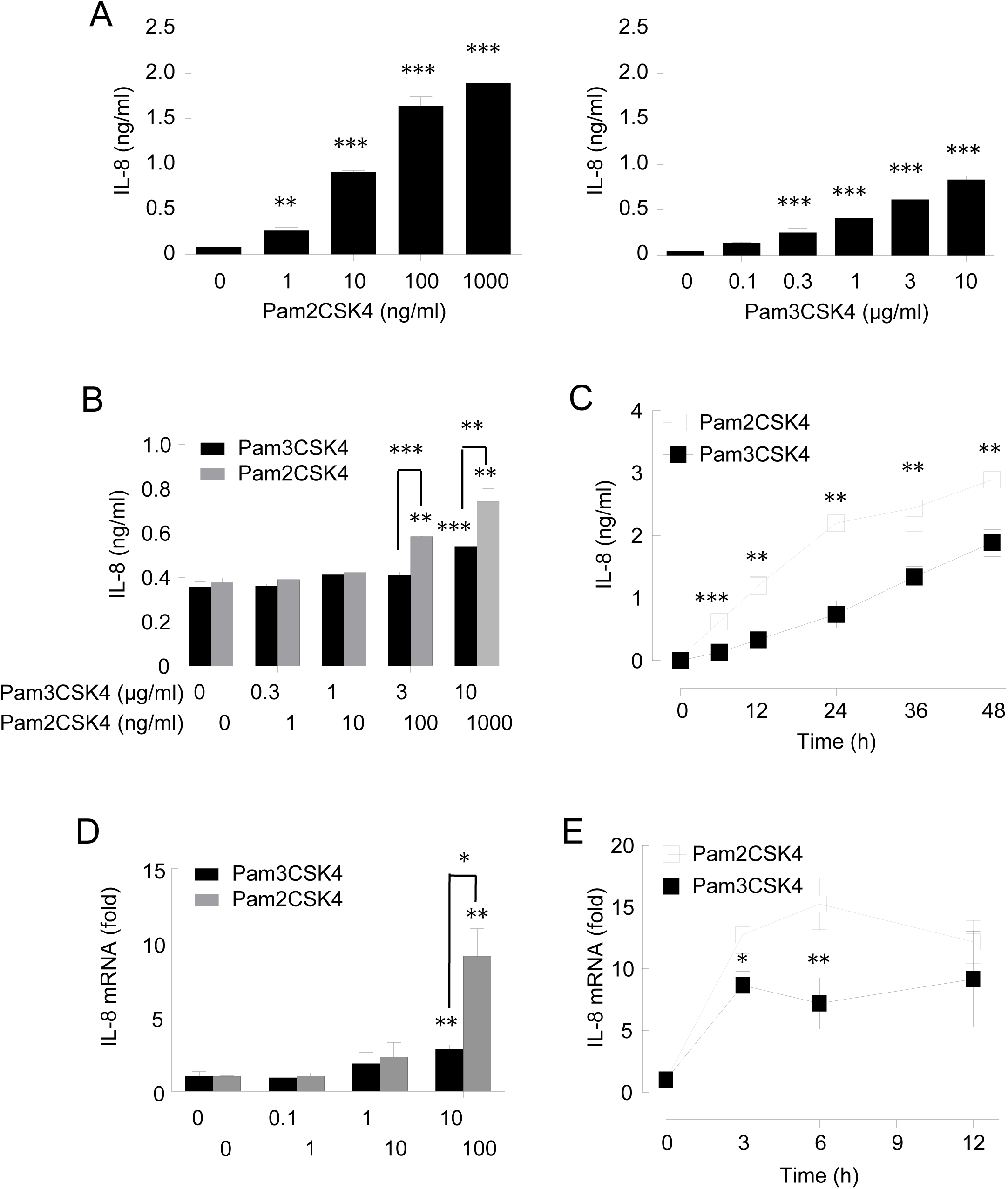
Diacylated and triacylated lipopeptides induce IL-8 expression with different potencies. (A) Synthetic lipopeptides pam2CSK4 (diacylated) and pam3CSK4 (triacylated) were treated on A549 cells in various concentrations for 24 h. IL-8 secretion was determined by ELISA. (B) IL-8 secretion in human primary bronchial epithelial cells stimulated with lipopeptides in various concentrations. Cells were starved for 6 h and then treated with lipopeptides for 24 h. (C) Time-dependent IL-8 secretion in cells treated with lipopeptide, pam2CSK4 (100 ng/ml) or pam3CSK4 (10 μg/ml). (D) IL-8 mRNA expression in A549 cells stimulated with lipopeptides in various concentrations for 24 h. IL-8 mRNA expression was measured by real-time qRT-PCR. (E) Time-dependent IL-8 mRNA expression upon treatment with pam2CSK4 (100 ng/ml) or pam3CSK4 (10 μg/ml). These suboptimal lipopeptide concentrations were used in following experiments, otherwise noted. Bars indicate means ± S.D. (n=3)

### Lipopeptides signal through intracellular TLR2 in A549 cells

Lipoproteins are preferentially recognized by TLR2 (Thorley et al., 2011). Indeed, transfection of TLR2 siRNA inhibited IL-8 secretion in both pam2CSK4- and pam3CSK4-treated cells (Fig. 3A). The siRNA reduced TLR2 mRNA expression by 80% (Fig. S1). We validated a role of TLR2 by blocking cell surface TLR2, CD14 or TLR4 in parallel using functional antibodies specific to each antigen. Interestingly, none of the antibodies tested affected on IL-8 secretion induced by lipopeptides (Fig 3B). This discrepancy between knockdown and blocking antibody studies suggested that functional TLR2 might reside in intracellular compartments. We examined this possibility using immunofluorescence. Cells were immunostained with fluorophore-conjugated TLR2-specific antibody in the presence or absence of cell-permeabilizing reagents and presence of TLR2 was examined using flowcytometry. Indeed, only permeabilized cells exhibited TLR2 fluorescence while intact cells did not show any significant fluorescence (Fig 3C). The result indicated that lipopeptides induce IL-8 expression via activating intracellular TLR2.

**Fig. 3.**
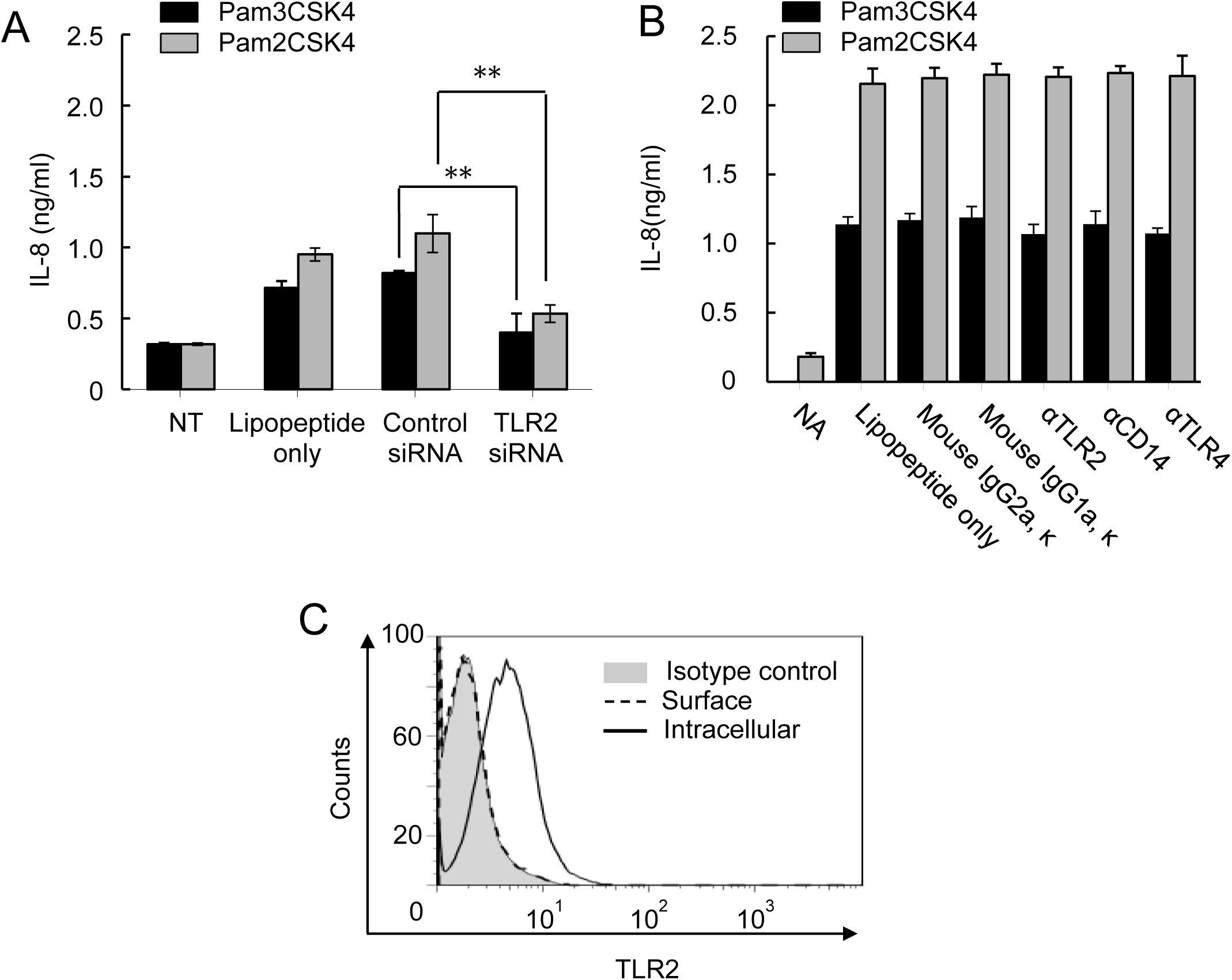
Lipopeptides induce IL-8 expression via intracellular TLR2 in A549 cells. (A) TLR2 knockdown effect on IL-8 production induced by lipopeptides in A549 cells. siRNA-transfected cells were treated with either pam2CSK4 or pam3CSK4 for additional 24 h. (B) Blocking antibodies for TLR2, TLR4, and CD14 had no effect on lipopeptide-induced IL-8 production. Cells were treated with blocking antibodies for 1 h prior to the lipopeptide stimulation for 24 h. Mouse IgG2aκ (for TLR2 and TLR4 antibodies) and IgG1aκ (for CD14 antibody) were used as isotype controls. Bars indicate means ± S.D. (n=3). (C) Intracellular localization of TLR2 in A549 cells was determined by immunostaining and FACS. Grey areas indicate isotype control. Dotted lines indicate fixed cells for surface staining and closed lines indicate permeabilized cells for intracellular staining of TLR.

### Pam2CSK4 and pam3CSK4 utilize different internalization mechanisms to activate TLR2 signaling

We hypothesized that lipopeptides might utilize an internalization mechanism to activate the intracellular TLR2 in A549 cells. It has been reported that PAMP-induced signal transduction requires ligand endocytosis (Brandt, Fickentscher, Kruithof, & de Moerloose, 2013; Bu, Wang, Tang, Koti, & Tan, 2010; Nilsen et al., 2008). For example, LTA internalizes via lipid-raft microdomain in the plasma membrane (Triantafilou et al., 2004). We tested different pharmacological inhibitors on IL-8 secretion to block potential ligand internalization mechanism. First, a role of lipid raft was examined by cholesterol extraction reagent methyl-beta-cyclodextrin (MβCD). MβCD pretreatment reduced pam2CSK4-induced IL-8 secretion dose-dependently by 25 % while it did not affect pam3CSK4-induced IL-8 secretion (Fig. 4A). The decrease was not a result from impaired cell viability (Fig. S2). Next, a role of clathrin-mediated endocytosis (CME) was examined using chlorpromazine that inhibits CME (Hussain, Leong, Ng, & Chu, 2011). Chlorpromazine did not affect pam2CSK4-induced response whereas it increased pam3CSK4-induced response (Fig. 4B). Finally, a role of macropinocytosis was examined using cytochalasin D that inhibits actin polymerization (Barthwal et al., 2013). Lipopeptide-induced IL-8 secretion was not significantly affected by this treatment in both pam2CSK- and pam3CSK4-treated cells. (Figure 4C). These results suggest that pam2CSK4 utilizes a lipid-raft dependent pathway whereas pam3CSK4 utilizes a CME-dependent pathway.

**Fig. 4.**
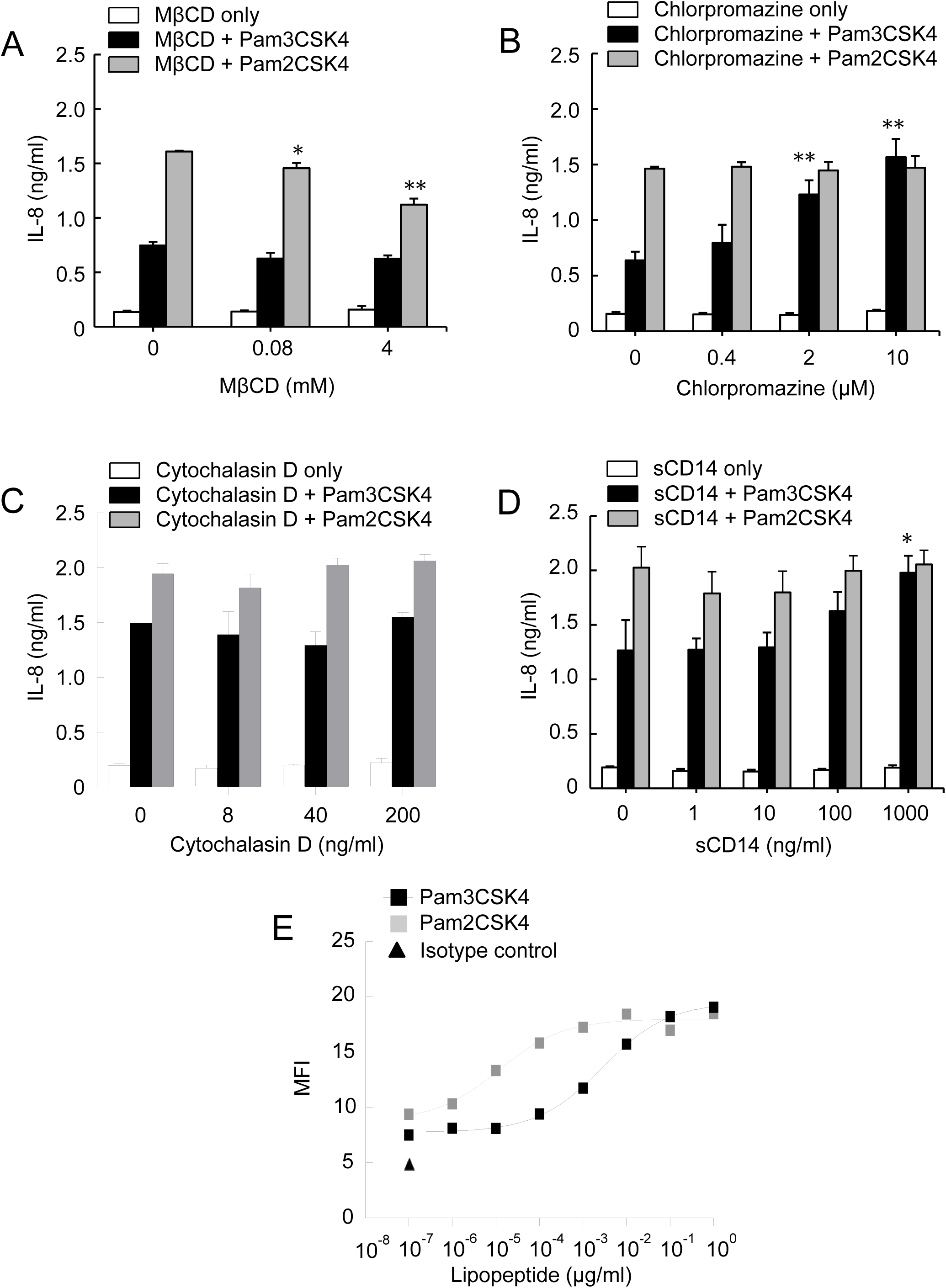
Lipopeptides internalize signals with distinct mechanisms. (A) A role of cholesterol-enriched lipid-raft in lipopeptide-induced IL-8 secretion in A549 cells. (B) A role of clathrin-mediated endocytosis in lipopeptide-induced IL-8 secretion. A549 cells were pre-treated with either MβCD or chlorpromazine for 1 h in indicated concentration and then stimulated with lipopeptides. (C) A role of macropinocytosis in lipopeptide-induced IL-8 secretion in A549 cells. Cells were pre-treated with various concentrations of cytochalasin D for 1 h and then lipopeptides-containing media were added for stimulation. (D) An effect of soluble CD14 (sCD14) on lipopeptide-induced IL-8 secretion. sCD14 were treated for 1 h prior to lipopeptide treatment (100 ng/ml Pam2CSK4 or 10 μg/ml Pam3CSK4). IL-8 secretion in the supernatant was determined by ELISA. Bars indicate means ± S.D. (n=3). (E) TLR2 signaling activity in CHO/CD14/TLR2 cells upon lipopeptide stimulation. Cells were treated with indicated concentrations of pam2CSK4 or pam3CSK4 for 24 h and then analyzed by flow cytometry for CD25 expression. Mean fluorescence intensity (MFI) is shown.

Chlorpromazine study suggested presence of a cell surface receptor that accumulates on the plasma membrane upon CME inhibition. This could be CD14 as it facilitates pam3CSK4-induced TLR2 activation (Nakata et al., 2006). For this reason, soluble CD14 (sCD14) was added prior to lipopeptide incubation since A549 cells do not exhibit cell surface CD14 (Thorley et al., 2011). Indeed, sCD14 augmented the pam3CSK4-induced IL-8 secretion to a level similar as pam2CSK4-induced IL-8 secretion but had no additive effect on pam2CSK4-induced response (Fig. 4D). This result was supportive of the hypothesis that pam3CSK4 might utilize CD14 to activate intracellular TLR2. Collectively, these data indicated that pam2CSK4 and pam3CSK4 utilize distinct endocytic pathways to activate intracellular TLR2.

Lipid structure of PAMP is important for TLR2 recognition (Hong et al., 2014) and therefore, different acylation in lipopeptides could affect TLR2 activation. We tested this hypothesis using CHO/CD14/TLR2 cells that reconstitute TLR2 signaling pathway in non-immune cells (Schroder et al., 2004). Cells were treated with varying concentrations of pam2CSK4 or pam3CSK4 and TLR2 signaling activity was monitored by cell surface expression of CD25 and flowcytometry (Fig. 4E). Both pam2CSK4 and pam3CSK4 increased CD25 expression in a dose-dependent manner. However, pam2CSK4 activated TLR2 signaling with higher efficiency than that of pam3CSK4. EC50 of pam2CSK4 and pam3CSK4 was 0.01 ng/ml and 3 ng/ml, respectively, indicating a 300-fold higher efficiency for pam2CSK4 than pam3CSK4. This experiment indicated that lipid moiety of lipopeptides determined potency of TLR2 activation.

### Pam2CSK4 and pam3CSK4 require different MAPK pathways for IL-8 induction

MAPK pathways regulate IL-8 secretion (Bhattacharyya et al., 2011). Involvement of MAPK activation was examined for lipopeptide-induced IL-8 secretion using pharmacological inhibitors for ERK1/2, p38 and JNK. An inhibitor of ERK1/2, U0126 preferentially reduced pam3CSK4-induced IL-8 secretion in a dose-dependent manner by 40% of control at 40 μM, while it had a minimal effect on pam2CSK4-induced response (Fig. 5A). The reduction did not result from impaired cell viability since U0126 did not affect cell viability at concentrations we used (Fig. S3A). A similar result was obtained when another ERK1/2 inhibitor with different chemistry PD98059 was tested (Fig. S3B). A role of p38 and JNK was examined in a similar manner. Interestingly, the p38-specific inhibitor SB203508 preferentially reduced pam2CSK4-induced IL-8 secretion by 60 % at 20 μM, with no significant effect on pam3CSK4-induced IL-8 (Fig. 5B). SB203508 did not significantly impair cell viability within concentration tested (Fig. S4A). Similar results were obtained when another p38 inhibitor SB202190 was used (Fig. S4B). JNK-specific inhibitor SP600125 slightly reduced pam2CSK4-induced IL-8 secretion while pam3CSK4-induced response was not affected (Fig. 5C). However, cell viability was slightly impaired in the highest dose of JNK inhibitor (> 10 μM) (Fig. S4C). Within lower concentration range that did not affect cell viability, SP600125 slightly reduced pam2CSK4-induced IL-8 secretion. The results showed that pam3CSK4 utilizes ERK1/2 pathway while pam2CSK4 utilizes p38 and JNK pathways.

**Fig. 5.**
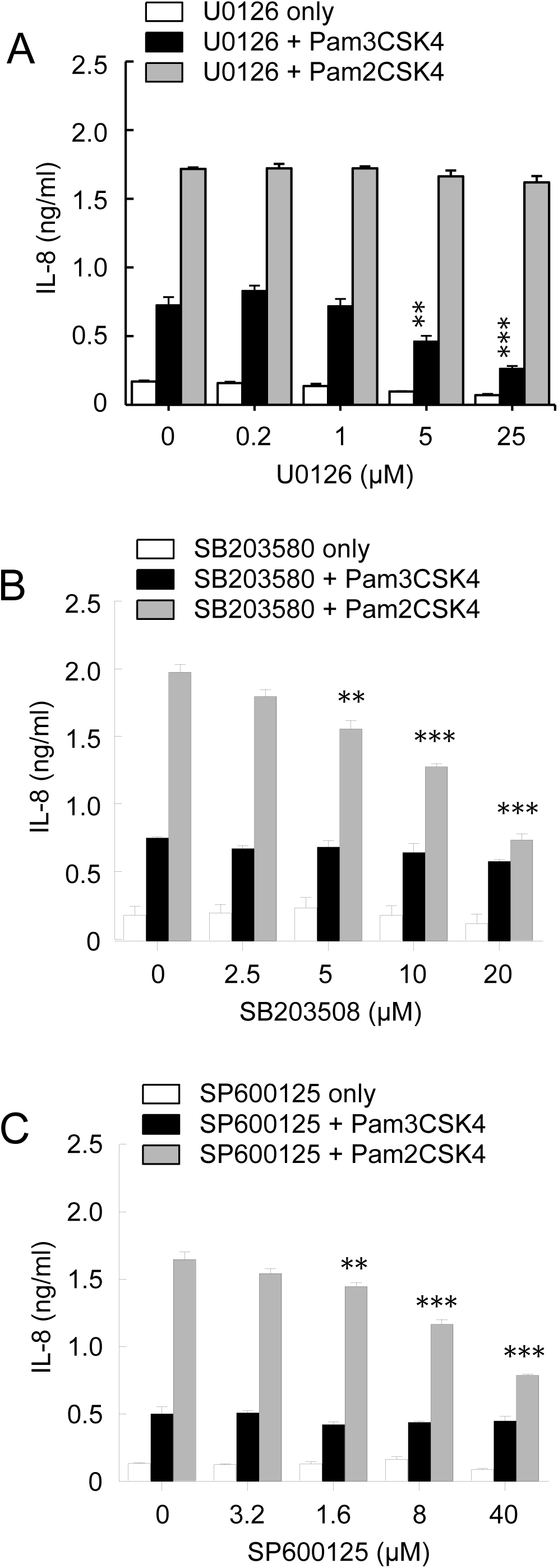
Lipopeptides differ in MAPK activation. A role of MAP kinases in lipopeptides-induced IL-8 production in A549 cells. Cells were pre-treated with various concentration of specific inhibitors for (A) ERK1/2 (U0126), (B) p38 kinase (SB203508) or (C) JNK/SAPK (SP600125) for 1 h and then stimulated with lipopeptides. The culture supernatants were analyzed for IL-8 secretion by ELISA technique. Bars indicate means ± S.D. (n=3).

### AP-1 and NF-κB are responsible for the higher IL-8 expression by pam2CSK4

IL-8 gene promoter contains *cis*-acting elements for transcription factors AP-1, NF-κB and NF-IL6 (Hoffmann, Dittrich-Breiholz, Holtmann, & Kracht, 2002; Jundi & Greene, 2015; Mukaida, Okamoto, Ishikawa, & Matsushima, 1994). Since pam2CSK4 and pam3CSK4 induce IL-8 gene expression with distinct mechanism, we reasoned that their capability to activate transcription factors for IL-8 expression might differ. For this, electromobility shift assay (EMSA) was utilized to determine transcription factor activation. Radio-labeled DNA probes containing consensus sequences for AP-1, NF-IL6 (alternatively called C/EBPβ), and NF-κB showed mobility shift when mixed with nuclear extract from pam2CSK4 or pam3CSK4-treated cells. Specific binding was tested by mixing with unlabeled probes. Both pam3CSK4 and pam2CSK4 increased band intensity of AP-1 probe- (Fig. 6A top panel) and NF-IL6 probe-protein complexes (middle panel). The most dramatic increase occurred for NF-κB probe-protein complex with 10-fold and 21-fold increase in band intensity for pam3CSK4-treated and pam2CSK4-treated samples, respectively (Fig. 6A bottom panel). These results indicate that lipopeptides increases DNA binding activity of the transcription factors that regulate IL-8 mRNA expression, with stronger activities of them in pam2CSK4-treated samples than in pam3CSK4-treated samples.

**Fig. 6.**
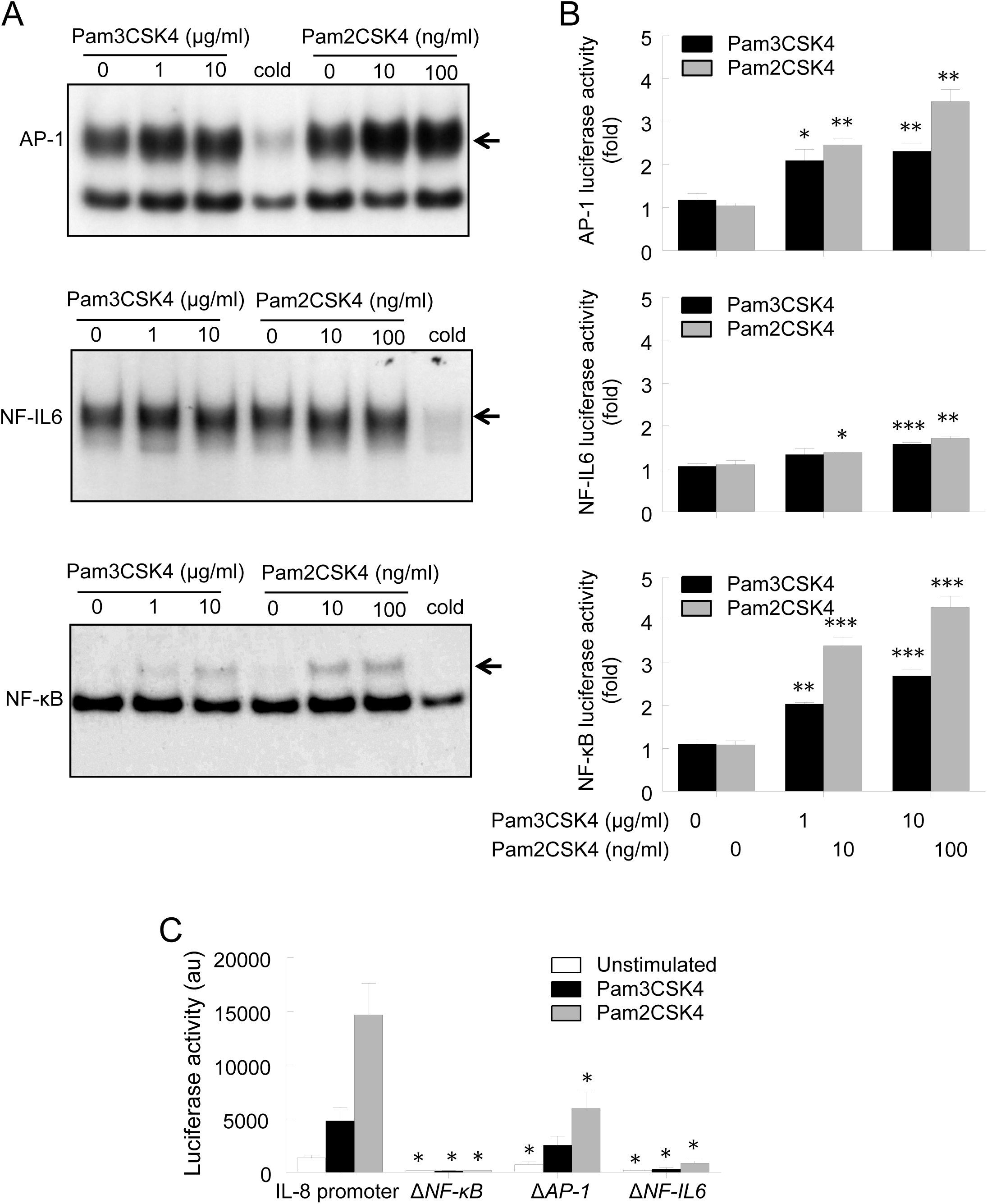
Lipopeptides differ in transcription factor activation for IL-8 expression. (A) Transcription factor activation upon lipopeptide stimulation was measured by their DNA binding activities using EMSA technique. DNA probes containing consensus sequence for AP-1, NF-IL6, and NF-κB binding were mixed with A549 cell nuclear extract from cells treated with lipopeptides. Images show autoradiographs for ^32^P-labled DNA probe-transcription factor complex. Specific bands were determined by competition with unlabeled DNA probes against the ^32^P-labled DNA-transcription factor complex (Cold). The results shown are representative of three different experiments. (B) Transcription factor activity upon lipopeptide stimulation using luciferase assay. Cells were transiently (24 h) transfected with luciferase plasmids containing consensus sequence for, AP-1, NF-IL6, or NF-κB. *Renilla* luciferase construct was co-transfected to detect transfection efficiency. Transfected cells were treated with lipopeptides for an additional 24 h. Data were normalized to unstimulated control. (C) Transcription factor requirement for IL-8 transcription upon lipopeptide treatment. Cells were transiently transfected with pGL3 plasmids containing either native human IL-8 promoter (WT) or mutant promoters with site-specific deletion of NF-κB, AP-1 or NF-IL6 binding site. Transfected cells were stimulated with either pam2CSK4 (100 ng/ml) or pam3CSK (10 μg/ml) for 24 h and cell lysates were subjected to luciferase assay. Difference in transfection efficiency was normalized by co-transfected *Renilla* luciferase activity. Bars indicate means ± S.D. (n=3).

We then examined activity of individual *cis*-acting elements from IL-8 promoter using luciferase reporter constructs (Fig. 6B). Both pam3CSK4 and pam2CSK4 treatments showed slight but significant increases (2 to 4 folds) in luciferase expression for all reporter constructs tested. Even in this case, pam2CSK4 consistently exhibited stronger activity than pam3CSK4 for luciferase expression. Native IL-8 promoter (−132 to +41 bp) showed much greater luciferase expression upon lipopeptide treatment. The role of individual *cis*-acting elements were validated by site-directed mutagenesis for each transcription factor binding sites (Im et al., 2009). The native IL-8 promoter construct (WT) robustly increased luciferase activity upon lipopeptide stimulation and the stronger pam2CSK4 response was consistent (Fig. 6C). Deletion of NF-κB and NF-IL6 virtually eliminated IL-8 promoter activity in both pam3CSK4- and pam2CSK4-stimulated cells indicating that NF-κB and NF-IL6 activation are crucial for lipopeptide-induced IL-8 expression. AP-1 deletion reduced luciferase expression significantly by 50 % in pam2CSK4-treated cells while it did not affect pam3CSK4-treated cells significantly with little difference from WT. Therefore, AP-1 only in pam2CSK4-treated cells plays a role to facilitate IL-8 expression. The result was correlated with distance of each *cis*-acting element from TATA box in IL-8 promoter since the closest NF-κB playing the most crucial role (Jundi & Greene, 2015). Overall, EMSA and promoter activity analyses showed that NF-κB and NF-IL6 activation is required for IL-8 expression upon lipopeptide challenge but AP-1 might play more important role for IL-8 expression induced by pam2CSK4.

## Discussion

Pulmonary epithelial cells can provide an innate immune defense against bacterial infection by secreting cytokine and chemokine to recruit immune cells like neutrophils (Cheon et al., 2008; Im et al., 2009; Thorley et al., 2011). Nonetheless, their role in host defense is not extensively studied compared to professional immune cells like dendritic cells and macrophages. Particularly, identity of PAMPs and degree of their contribution to local inflammation have not been extensively studied. We previously showed that bacterial flagellin and PGNs induces IL-8 secretion in A549 cells while a role of other PAMPs remained unclear (Cheon et al., 2008; Im et al., 2009). In the current study, we report that diacylated lipoprotein from Gram-positive bacteria are a potent inducer of IL-8 secretion.

We found that lipoprotein from *S. aureus* induces the most potent IL-8 secretion response when compared with other Gram-negative and Gram-positive cell wall components (Fig. 1A). This was confirmed using lipoprotein-lacking mutant *S. aureus* strains (Fig. 1B). Surprisingly, synthetic lipopeptides with diacylated moiety (pam2CSK4) consistently exhibited higher IL-8 inducing capability throughout experiments using the human lung epithelial A549 cells and primary human bronchial epithelial cells than triacylated pam3CSK4. Diacylated lipoproteins are more abundant in Gram-positive bacteria as lipoprotein synthesis in Gram-positive bacteria occurs through two step reaction through lipoprotein diacylglyceryl transferase (Lgt) and lipoprotein signal peptidase (Lsp) while that of Gram-negative bacteria includes additional acylation step using lipoprotein N-acyl transferase (Lnt) (Kovacs-Simon et al., 2011). Therefore, data in our current study suggest that Gram-positive bacteria in lung microbiota would play a primary role in inducing epithelial immunity at least in part for IL-8 induction.

We suggest that there are multiple layers of important signal transduction checkpoints that result in the stronger pam2CSK4-induced IL-8 secretion: 1) pam2CSK4 was more efficient than pam3CSK4 in activating TLR2 regardless of whether there is TLR1 or TLR6 available; 2) pam2CSK4 utilized p38 and JNK pathways while pam3CSK4 preferentially utilized ERK1/2 pathway; 3) pam2CSK4 was able to engage AP-1 in IL-8 expression while pam3CSK4 did not. The difference we observed here between the lipopeptides was not due to their difference in molarity since, at the same concentration, molar ratio of pam3CSK4 to pam2CSK4 was 0.85, which does not correspond to the big difference we found from CHO/TLR2/CD14 cell study (Fig. 4E). Rather, the different activities more likely occurred via higher affinity of pam2CSK4 to TLR2. The presence of TLR1 or TLR6 might provide with more sophisticated regulation of signal transduction. A comparison between TLR2/TLR1 and TLR2/TLR6 complexes in 3D structure showed that triacylated lipoproteins require TLR1 to accommodate additional N-acyl group while TLR6 only confers more hydrophobic interface to diacylated lipoproteins (Kang et al., 2009). Therefore, TLR1 is likely important for pam3CSK4 recognition while TLR6 might be somewhat dispensable for pam2CSK4 recognition. Presumably, triacylated lipopeptides in general require a co-receptor to activate TLR2 such as TLR1 or CD14 (Fig. 4B, D). Since TLR2 in A549 cell localizes to intracellular compartments (Fig. 3), we reason that A549 cells might utilize a signaling pathway that is distinct from conventional MyD88-dependent pathway. It is known that MyD88 is in general required for lipoprotein-dependent immune response in *S. aureus* infection (Takeuchi, Hoshino, & Akira, 2000). However, information regarding its role for release of IL-8 in A549 cells is scarce. An adaptor other than MyD88 might play a role in this case provided that intracellular TLRs tend to switch to different adaptor like TRIF for downstream signaling (Barton & Kagan, 2009).

Differential MAPK activation might be controlled at the plasma membrane level. Disruption of lipid-raft is known to abolish p38 phosphorylation, but has no effect on ERK phosphorylation in T cell signaling (El Fakhry, Alturaihi, Diallo, Merhi, & Mourad, 2010). Likewise, cholesterol sequestration in lipid-raft reduced p38 and JNK activity in class A macrophage scavenger receptor signaling while down-regulation of clathrin inhibited ERK1/2 activation (Zhu et al., 2011). These reports concur with our results that different ligand internalization mechanisms result in distinct signal outcome, where lipid-raft dependent pathway activates p38 (pam2CSK4) while clathrin-mediated endocytic pathway activates ERK1/2 (pam3CSK4) (Figs 4, 5). p38 activation could bring additional induction of IL-8 secretion by stabilizing mRNA, which partly explains the higher IL-8 mRNA expression and secretion in pam2CSK4-treated cells (Fig. 2) (Holtmann et al., 1999). Therefore, distinct activation pattern we found for MAPKs and possibly transcription factor activation upon different lipopeptide treatment could be attributed to the upstream regulation at the plasma membrane.

Overall, our current study showed that lipid moiety in bacterial lipopeptides plays a crucial role in respiratory epithelial immunity. The results presented in the current work may provide a useful insight for pharmacological intervention point for acute or chronic lung inflammatory diseases that involve dysbiosis of local microbiome.

## Material and Methods

### Bacterial strains and reagents

Pam2CSK4 and pam3CSK4, PGNs from *S. aureus* and *E. coli*, and ultra-pure K12 LPS were purchased from InvivoGen (San Diego, CA, USA). Highly pure and structurally intact *S. aureus* LTA was purified by butanol extraction followed by hydrophobic column chromatography and ion-exchange column chromatography as previously described (Ryu et al., 2009). Ethanol-inactivated *S. aureus* RN4220, LTA-deficient mutant (Δ*ltaS*), and lipoprotein-deficient mutant (Δ*lgt*) were prepared as previously described (J. Kim et al., 2013). Lipoproteins from *S. aureus* and *E. coli* were isolated as previously described (A. R. Kim et al., 2017). Methly-β-cyclodextran (MβCD) was purchased from Sigma-Aldrich (St. Louis, MO, USA). All specific inhibitors for MAPKs were obtained from Calbiochem (La Jolla, CA, USA). Functional grade monoclonal antibodies for blocking TLR2 (TL2.1), CD14 (61D3) and TLR4 (HTA125) and their matching isotype antibodies were purchased from eBioscience (San Diego, CA, USA). CD25 antibody was purchased from BD Bioscience (clone 2A3).

### Cell culture

A549 cells were purchased from American Type Culture Collection (Manassas, VA, USA) and maintained in Ham’s F12 nutrient mixture containing 2 mM L-glutamine, 1.5 g/L sodium bicarbonate supplemented with 10% heat-inactivated fetal bovine serum and 1% streptomycin and penicillin (Life Technologies, Grand Island, NY, USA). Human primary bronchial epithelial cells were purchased from PromoCell (Heidelberg, Germany) and grown in an airway epithelial cell growth medium with airway epithelial cell supplement pack (PromoCell) according to the manufacturer’s instruction. CHO/CD14/TLR2 cells, which constitutively expresses human CD14 and TLR2, and expresses membrane-bound CD25 upon TLR2 activation, was cultured and used as previously described (Hong et al., 2014).

### Enzyme-linked immunosorbent assay (ELISA)

A549 cells (1 × 10^5^ cells/well) were seeded in 48 well cell culture plates 16 h prior to PAMP stimulation, which was typically performed by 24 h incubation. After stimulation, cell culture supernatants were collected and IL-8 secretion was determined using IL-8 ELISA kit (R&D Systems, Minneapolis, MN, USA) according to the manufacturer’s instructions. For inhibitor experiments, A549 cells were pre-treated either with specific blocking antibodies for TLR2, TLR4 and CD14 or with specific inhibitors against ERK1/2, p38 kinase and JNK/SAPK for 1 h followed by lipopeptide stimulation for an additional 24 h. The blocking antibodies and inhibitors were co-present with lipopeptides during the stimulation period. Following stimulation, culture supernatants were collected to determine IL-8 secretion by ELISA. Human primary bronchial epithelial cells (2 × 10^5^ cells/ml) were plated for 24 h followed by starvation in serum-free medium for 6 h. Then, the cells were stimulated in the same manner as A549 cells and level of IL-8 was determined as described above. Cell viability upon inhibitor treatment was measured by MTT assay as previously described (Cheon et al., 2008). In brief, MTT solution (0.5 mg/ml) was added to the cells after transferring supernatants for IL-8 measurement and incubated at 37 °C in 5 % CO_2_ for 40 min. After removal of media containing MTT solution, cells were dissolved in dimethylsulfoxide (DMSO) and color changes in cells was detected by absorbance at 550 nm using a microtiter plate reader (VersaMax, Molecular Devices, Sunnyvale, CA, USA).

### Real-time polymerase chain reaction (Real-time RT-PCR)

Total RNAs were extracted using TRIzol reagent (Invitrogen, Calsbad, CA, USA) and cDNAs were synthesized using M-MLV reverse-transcriptase (Promega, Madison, WI, USA) using random hexamer nucleotides as primers (Promega). Real-time RT-PCR for the cDNAs was performed using *TaqMan* enzyme mixture for manufacturer-designed human CXCL-8, human TLR2, and human GAPDH *TaqMan* probes conjugated with FAM6 dye which were purchased from Applied Biosystems (Cat# 4331182 Applied Biosystems, currently merged with Thermo Scientific). PCR reaction performed using ABI Prism 7500 Sequence Detection System (Applied Biosystems).

### Flow cytometric analysis

For intracellular TLR2 expression, A549 cells were permeabilized with Cytofix/Cytoperm Plus fixation/permeabilization kit (BD Biosciences, San Jose, CA, USA) and stained with FITC-conjugated anti-TLR2. Cells were stained with the anti-TLR2 antibody without permeabilization for surface TLR detection. TLR expression was analyzed by flowcytometry using a FACSCalibur flowcytometer operated with CellQuest software (BD Biosciences). For CHO/CD14/TLR2 cell activation, cells were stimulated with either pam2CSK4 or pam3CSK4 for 24 h and surface CD25 expression was analyzed using specific CD25 antibody as described above but without permeabilization.

### Transfection with small interfering RNA (siRNA)

A549 cells (3 × 10^4^ cells/well) were plated in 48-well culture plates for 24 h prior to transfection with human TLR2 siRNA (sc-4025, Santa Cruz) or non-targeting control siRNA (sc-37007, Santa Cruz, CA, USA) using Lipofectamine RNAiMAX reagent (Invitrogen) according to the manufacturer protocol. Six hours after the transfection, siRNA containing media were replaced with complete Ham’s F12 nutrient mixture and incubated for additional 24 h in the presence of pam2CSK4 or pam3CSK4. IL-8 production was measured by ELISA for the culture supernatants.

### Electrophoretic mobility shift assay (EMSA)

A549 cells (1 × 10^6^ cells/100 mm dish) were treated with either pam2CSK4 or pam3CSK4 for 90 min and nuclear extracts were prepared as follows. Briefly, the cells were lysed with hypotonic buffer (10 mM HEPES and 1.5 mM MgCl_2_) and subsequently nuclei were pelleted by centrifugation at 3000 rpm for 5 min and lysed in lysis buffer (30 mM HEPES, 1.5 mM MgCl_2_, 450 mM KCl, 0.3 mM EDTA, 10% glycerol, 1 mM DTT, 1 mM PMSF and 1 μg/ml aprotinin and 1 μg/ml leupeptin). After centrifugation at 15000 rpm, supernatants were kept at −80 °C until used. EMSA was performed with double-stranded deoxyoligonucleotide probes for NF-κB, AP-1 and NF-IL6 consensus sequences as previously described (Im et al., 2009).

### Transfection and luciferase reporter gene assay

A549 cells were transfected with NF-κB-Luc, AP-1-Luc or NF-IL6-Luc plasmid (Clontech, Pal Alto, CA, USA) using Lipofectamine 2000 for 24 h (Stratagene, La Jolla, CA, USA). For IL-8 native promoter experiment, cells were transfected with either WT construct containing native human IL-8 promoter/pGL3 or mutant/pGL3 constructs containing site-specific deletion for NF-κB, AP-1 or NF-IL6 transcription factor-binding site (Im et al., 2009). All luciferase constructs were co-transfected with pRL-TK *Renilla* luciferase plasmid (Promega, Madison, WI, USA) to normalize transfection efficiency variation between samples. Luciferase transfections were for 24 h. After the transfection, transfected cells were stimulated with pam2CSK4 or pam3CSK4 for an additional 24 h. Then, the cells were lysed with Glo lysis buffer (Promega) and luciferase activity was analyzed by using Victor 1420 Multi-label counter (PerkinElmer Life and Analytical Sciences, Waltham, MA, USA).

### Statistical analysis

Experiments were performed at least two times with triplication. Means ± standard deviation (S.D.) were calculated from triplicated samples, otherwise mentioned. Statistical significance was tested by Welch’s *t*-test. *, **, and *** indicate *p*<0.05, *p*<0.01 and *p*<0.001, respectively.

## Acknowledgements

This work was supported by National Research Foundation grants of South Korea (2010-0029116, 2008-0062421 to Dr. Seung Hyun Han, Seoul National University Dental School). I greatly appreciate the generous support for this work from the lab of Dr. Han.

---

**Fig. S1.**
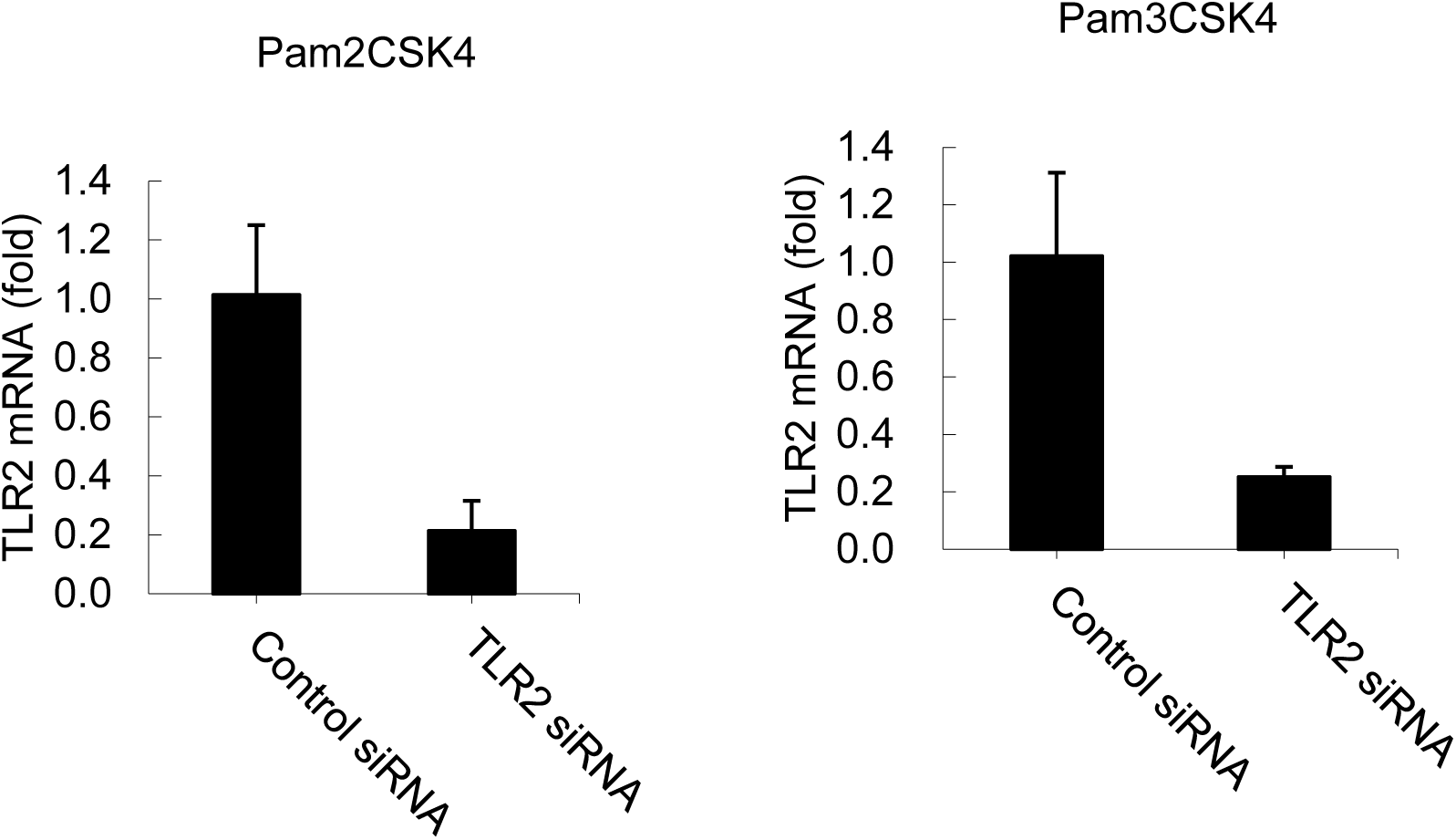
TLR2 knockdown efficiency. A549 cells were transfected with either control or TLR2 siRNA for 6 h and then treated with either pam2CSK4 or pam3CSK4 for additional 24 h to determine IL-8 secretion. The cells were collected and TLR2 mRNA expression was determined by quantitative RT-PCR from total RNA. Bars indicate means ± S.D. (n=3).

**Fig. S2.**
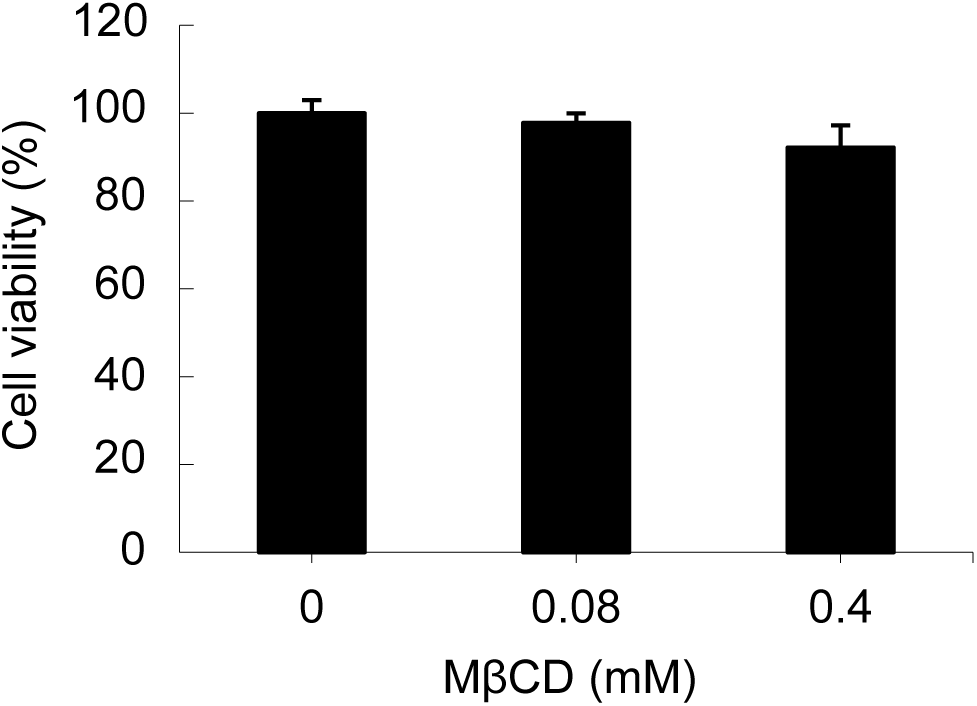
Effects of MβCD treatment on cell viability. A549 cells were incubated with various concentrations of MβCD for 24 h. Cell viability was determined by MTT assay. Bars indicate means ± S.D. (n=3).

**Fig. S3.**
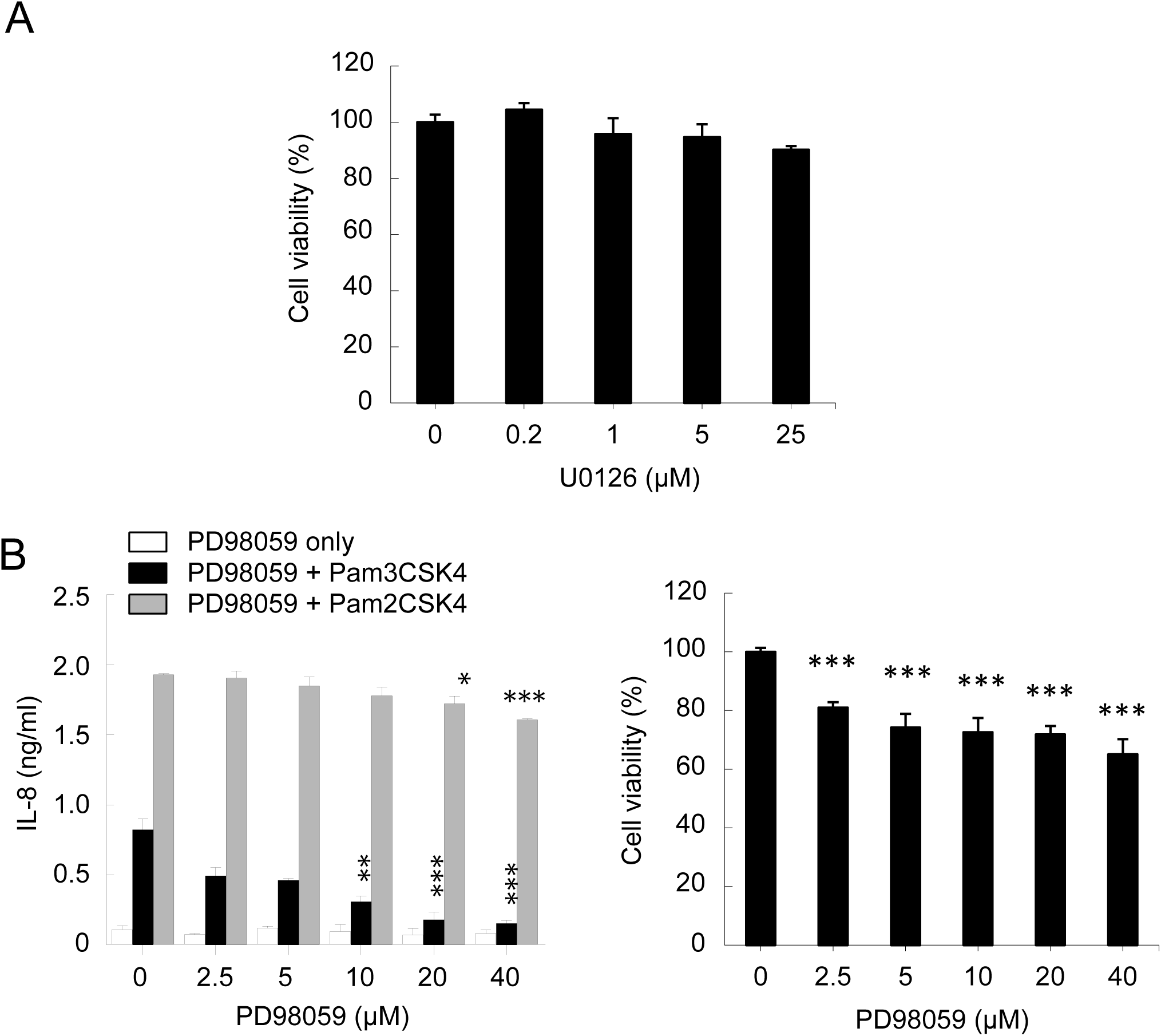
Effects of ERK1/2 inhibition on lipopeptides-induced IL-8 production. (A) A549 cell viability determined by MTT assay at 24 h after U0126 treatment. (B) Effect of PD98059 on lipopeptide-induced IL-8 secretion. Note that cell viability was reduced by ∼80% in all sample treated with PD98059 (right-side panel) except control. Pam3CSK4-induced IL-8 secretion was however reduced by ∼90% when compared with pam3CSK4 only control (0 μM; left-side panel). Therefore, the large reduction in pam3CSK4-induced IL-8 secretion was specifically inhibited by PD98059 whereas pam2CSK4-induced IL-8 secretion was relatively unaffected. Bars indicate means ± S.D. (n=3).

**Fig. S4.**
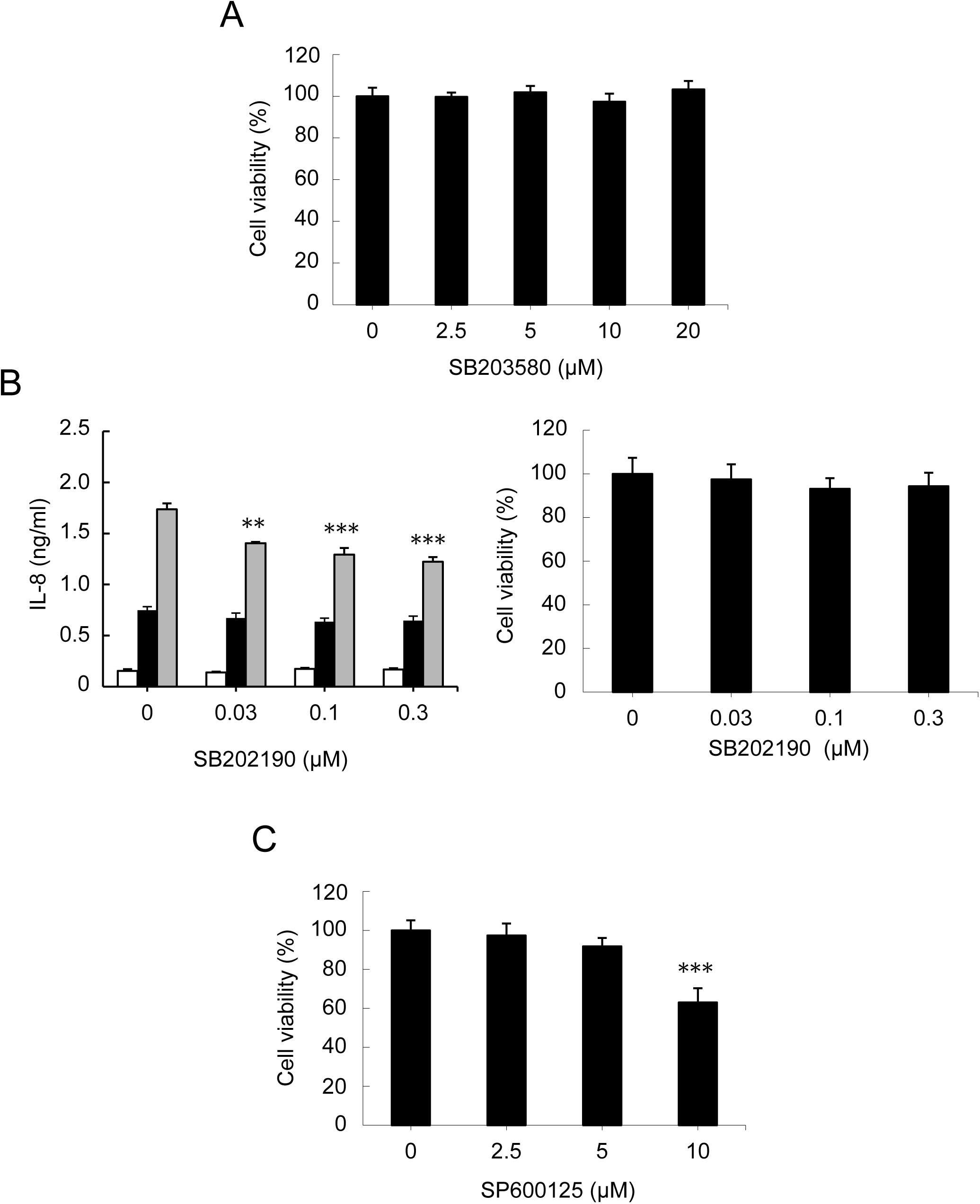
P38 and JNK inhibition in IL-8 induction. (A) Cell viability measured by MTT assay after SB203580 (p38 inhibitor) treatment for 24 h. (B) Effect of SB202190 on lipopeptide-induced IL-8 secretion in A549 cells. Cell viability remained unaffected by the inhibitor treatment (right-side panel). Pam2CSK4-induced IL-8 secretion was specifically reduced by pretreatment of SB202190 for 1 h. (C) Effect of JNK inhibitor SP600125 treatment on A549 cell viability. Cell viability was reduced by ∼30 % upon treatment of 10 μM SP600125 while cells remained unaffected in lower concentrations.

